# Molecular evolution of octopamine receptors in *Drosophila*

**DOI:** 10.1101/2025.10.30.685676

**Authors:** Mengye Yang, Jolie A. Carlisle, Ben R. Hopkins, Mariana F. Wolfner

## Abstract

Octopamine (OA), the insect analog of noradrenaline, plays important roles in diverse behavioral and physiological processes, from modulating fight-or-flight behavior to regulating post-mating ovulation. In *Drosophila*, six OA receptors have been identified: Oamb, Octα2R, Octβ1R, Octβ2R, Octβ3R, and Oct-TyrR, and they have been linked to different behavioral and physiological processes. Here, we investigated the evolutionary characteristics of these receptors across *Drosophila* species. We found that OA receptors are generally found as single-copy genes. Notably, Octβ2R and Octβ3R exhibit positive selection within the *melanogaster* group, though in different structural regions from one another. The positively selected sites in Octβ2R are exclusively located in regions important for ligand binding, whereas those in Octβ3R are predominantly found in regions crucial for signal transduction. Interestingly, Octβ2R remains highly conserved outside the *melanogaster* group, so the detection of positive selection in its ligand-binding related domains within this clade raises the possibility that it has evolved an additional, *melanogaster*-specific ligand interaction(s), among other potential reasons. These findings highlight the evolutionary flexibility of aminergic signaling and suggest lineage-specific adaptations of OA receptor function in *Drosophila*, likely shaped by lineage-specific selective pressures.

**Article summary:** Octopamine is a chemical messenger in insects, analogous to noradrenaline in vertebrates, and influences many processes such as fight-or-flight responses, locomotion, and reproduction. In fruit flies and other insects, multiple octopamine receptors detect and respond to this signal. We explored how these receptors have changed over millions of years across various fly species. We found that while some receptors remain similar, others have evolved more rapidly, possibly due to changes in the insects’ lifestyles, environments, or other biological demands. Understanding these differences can help reveal how insects fine-tune their behaviors and adapt to diverse challenges.

## Introduction

Biogenic amines are small signaling molecules that play multiple roles in regulating physiological and behavioral functions across both mammals and insects. In mammals, noradrenaline and adrenaline mediate fight-or-flight responses and other key processes. In insects, however, these catecholamines are absent or present only at very low levels. Instead, insects rely on structurally and functionally analogous molecules: tyramine (TA) and octopamine (OA). The amino acid tyrosine is first converted to TA by tyrosine decarboxylase, and TA is subsequently converted to OA by tyramine β-hydroxylase. Although TA serves as the biosynthetic precursor to OA, both can act as independent neurotransmitters (Roeder, 2005). In *Drosophila melanogaster* (fruit fly), OA function has been particularly well-studied, revealing its pleiotropic roles in diverse processes including locomotion, aggression, learning, memory, sleep, energy intake and expenditure, as well as reproduction (Farooqui, 2012; Roeder, 2005, 2020; White et al., 2021).

OA exerts its diverse physiological and behavioral effects by binding to and activating its receptors, all of which belong to the G protein-coupled receptor (GPCR) family. In *Drosophila*, these receptors are classified into four groups based on their structural and signaling similarities to vertebrate adrenergic receptors: Oamb (α-adrenergic-like octopamine receptor), Octα2R (α_2_-adrenergic-like octopamine receptor), three β-adrenergic-like receptors (Octβ1R, Octβ2R, and Octβ3R), and the octopamine/tyramine receptor Oct-TyrR (Evans & Maqueira, 2005; Qi et al.,2017). Despite being functionally related proteins belonging to the same protein family, the relationships among the OA receptors remains unresolved. Previously, two independent evolutionary studies constructed phylogenetic trees of biogenic amine receptors across species. As expected, β-adrenergic-like octopamine receptors clustered together; however, the phylogenetic and evolutionary relationships with and among the other OA receptors remained unclear due to weak branch support (Qi et al., 2017; Zhang et al., 2023). Upon activation, Oamb promotes increases in intracellular calcium and cAMP levels and has been implicated in processes such as follicle rupture, ovulation, sperm storage, sleep/wake regulation, appetitive learning and memory, insulin-like peptide transcription, and male aggression (Avila et al., 2012; Balfanz et al., 2005; Burke et al., 2012; Crocker et al., 2010; Deady & Sun, 2015; Han et al., 1998; Huetteroth et al., 2015; Kim et al., 2013; Lee et al., 2009; Luo et al., 2014). Octα2R, by contrast, reduces cAMP production and has been linked to locomotion, grooming behavior, and starvation-induced hyperactivity (Nakagawa et al., 2022; Qi et al., 2017). The OctβRs predominantly signal through cAMP, and Octβ1R is involved in olfactory learning, exercise adaptation, and hunger-driven modulation of female receptivity (Sabandal et al., 2020; Sujkowski et al., 2020; Sun et al., 2023); Octβ2R regulates ovulation, locomotor activity, anesthesia-resistant memory, and sleep, as well as stimulates synaptic growth, an effect antagonized by Octβ1R (Koon et al., 2011; Koon & Budnik, 2012; Li et al., 2015; Lim et al., 2014; Wu et al., 2013; Zhao et al., 2021); while Octβ3R has been implicated in metamorphosis and appetitive motivation (Ohhara et al., 2015; Zhang et al., 2013). Finally, Oct-TyrR is sensitive to both TA and OA, with TA being slightly more potent in inhibiting adenylate cyclase activity, while OA more strongly stimulates calcium signaling (Robb et al., 1994). Oct-TyrR has been shown to affect chemotaxis behavior and startle responses by modulating downstream dopaminergic neuron activity (Ma et al., 2016).

Genes associated with reproduction not infrequently evolve more rapidly than non-reproductive genes, sometimes exhibiting elevated sequence divergence potentially driven by sexual selection, sexual conflict, or relaxed selective constraints (Carlisle & Swanson, 2021; Clark et al., 2006; Dapper & Wade, 2020; Swanson & Vacquier, 2002; Wilburn & Swanson, 2016). Male seminal fluid proteins trigger the female post-mating response, a potential battleground of male x female interaction. Indeed, 7-12% of male seminal fluid proteins show signatures of positive selection potentially driven by sexual selection or sexual conflict (Haerty et al., 2007; Patlar et al., 2021; Swanson, Clark, et al., 2001; Wong & Wolfner, 2012). Although fewer studies have addressed the evolution of female reproductive proteins, growing evidence suggests that positive selection also acts on the female side (Galindo et al., 2003; McGeary & Findlay, 2020; Moyle et al., 2021; Swanson et al., 2001). Given that multiple OA receptors are involved in reproductive processes (Avila et al., 2012; Deady & Sun, 2015; Lee et al., 2009; Li et al., 2015; Lim et al., 2014; Sun et al., 2023), it is of particular interest to explore the evolutionary dynamics of this receptor group.

Here, we identified OA receptor orthologs across 27 *Drosophila* species and detected two lineage-specific tandem duplication events using synteny analysis. Nevertheless, OA receptors are generally maintained as single-copy genes. Using a suite of molecular evolution analyses, we found evidence of positive selection on Octβ2R and Octβ3R within the *melanogaster* group.Interestingly, we did not detect evidence for positive selection of Octβ2R in the *virilis*-*repleta* radiation nor in mosquitoes, suggesting that the positive selection observed in residues important for ligand binding within *melanogaster* species may reflect lineage-specific selective pressures. These selective pressures could include sexual selection on reproductive function, pathogen-mediated selection, ecological adaptation involving receptor modulation, or perhaps co-option to bind a ligand other than OA. Taken together, these results reveal distinct evolutionary trajectories shaping OA receptor evolution.

## Materials and Methods

### Synteny analysis and tandem duplication detection

To identify potential tandem duplications of OA receptors, we first determined their syntenic regions by focusing on the three closest upstream and downstream genes flanking each OA receptor in the *Drosophila melanogaster* genome, as identified using the JBrowse map within FlyBase (flybase.org). Using three neighboring genes provides sufficient local context to detect tandem duplications without extending into unrelated genomic regions. We then identified orthologs of these seven genes (including the focal OA receptor) in 27 *Drosophila* species whose genomes had been annotated by the automated NCBI Gnomon prediction pipeline. The *D. melanogaster* sequence of each gene was used as the query in the tBLASTn search against the annotated transcriptomes for each species. The species we tested were: *melanogaster, simulans, mauritiana, sechellia, erecta, yakuba, santomea, teissieri, takahashii, suzukii, subpulchrella, biarmipes, elegans, rhopaloa, ficusphila, kikkawai, ananassae, persimilis, pseudoobscura, willistoni, busckii, grimshawi, mojavensis, hydei, virilis, innubila*, and *nasuta*. We also performed reciprocal tBLASTn searches against *D. melanogaster* transcriptome to strengthen the confidence in ortholog prediction. Their chromosomal locations were confirmed using the NCBI Genome Data Viewer, and OA receptor copy number was determined for each species. For species with more than one copy of an OA receptor, RNA-seq exon coverage tracks were inspected on the species’ genome browsers to confirm transcription of each identified paralog.For OA receptors, at least one side of the flanking genes showed some level of conservation, supporting their placement within the expected genomic context. For flanking genes that differed from those in *D. melanogaster*, we used Gnomon predictions available on NCBI to confirm their identities. Synteny metrics are provided in File S1, and more details are available in File S6.

High-throughput methods such as those used in OrthoDB (Tegenfeldt et al., 2025) and DROSOMA (Thiébaut et al., 2024) provide large-scale resources for ortholog annotation, but they rely on automated pipelines that can misclassify relationships, particularly in cases of gene duplication or lineage-specific loss (Carlisle et al., 2024). Our approach involves nonautomated, detailed, and targeted characterization of receptor genes across species. While our results largely agree with OrthoDB and DROSOMA for most octopamine receptors existing as single-copy in species examined, these databases did not reliably find the duplicates that we detected. The duplicate in *D. busckii* of *Octβ3R* was not detected in either DROSOMA or OrthoDB and the duplicate in *D. sechellia* of *Octα2R* was not detected in DROSOMA but was detected in OrthoDB. This inconsistency between our results and ortholog predictors and the inconsistency between ortholog predictors themselves highlight the value of our detailed, targeted investigations, such as that presented here.

### Molecular evolution analyses

We performed PAML analyses on each OA receptor across 15 species within the *melanogaster* group, as listed in Fig S4. Only isoforms detectable in more than 10 species by Gnomon prediction were included in the analysis. For Octα2R, *Dsec* orthologs were excluded due to the presence of two copies in this species. Protein sequence alignments were generated by Clustal Omega (Madeira et al., 2024), which were then converted into codon-based DNA alignments with PAL2NAL (Suyama et al., 2006). Protein alignment statistics can be found in File S7. Maximum likelihood trees were constructed with reference to two established phylogenies: a high-confidence phylogeny of 155 *Drosophila* species (Suvorov et al., 2022) and an additional tree including *D. santomea* (Hopkins et al., 2024) to determine its phylogenetic position. The codeml program from the PAML package (Yang, 2007) was used to calculate an overall ω estimate for the whole sequence under model M0 and to perform site tests by comparing model M8, which allows for a class of sites with ω >1, against the null models M7 and M8a using likelihood ratio tests (Swanson et al., 2003; Yang et al., 2000). The ‘cleandata’ option was enabled in codeml to remove alignment sites with gaps or ambiguous data. For genes where model M8 provided a significantly better fit than models M7 and M8a, the BEB approach was applied to identify positively selected sites at a 0.9 confidence level.

For Octβ2R, PAML analyses were also performed within the *Drosophila virilis*-*repleta* radiation and across mosquito species, following the similar procedures described above. Ortholog prediction was conducted in the same manner as for the 27 *Drosophila* species (see the Synteny analysis and tandem duplication detection section), and PAML analyses were carried out as described in the preceding paragraph. Within the *virilis*-*repleta* radiation, seven species (*virilis, novamexicana, hydei, navojoa, mojavensis, arizonae*, and *montana*) were chosen based on the availability of transcriptome data. In mosquitoes, PAML tests were performed separately within the *Cellia, Anophelinae*, and *Culicidae* lineages. The species included in these analyses, along with their phylogenetic relationships (Neafsey et al., 2015; Suvorov et al., 2022), are shown in Fig 4B.

MEME (Mixed Effects Model of Evolution) analyses were conducted using the Datamonkey Adaptive Evolution Server (Weaver et al., 2018) with default parameters and 100 resamples. The maximum likelihood trees and DNA alignments used in the PAML analyses were also used as input for these analyses.

### Codon usage bias and GC content analysis

CDS of Octβ2R and Octβ3R from 15 species in the *Drosophila melanogaster* group were analyzed to assess codon usage bias and GC content. Sequences were first cleaned by removing gaps (-). Codon usage metrics were calculated using the coRdon R package (v1.20.0), including the effective number of codons (ENC) to quantify overall codon usage bias. Overall GC content and GC content in the third codon position (GC3) were computed using the seqinr R package (v 4.2.36). For GC3, the third nucleotide of each codon was extracted and the proportion of G or C nucleotides was calculated per sequence.

ENC values range from 20 (extreme codon bias) to 61 (no bias), with intermediate values indicating moderate codon usage bias. GC and GC3 metrics were used to evaluate potential nucleotide composition shifts that might influence ω estimates. All analyses were performed in R (v4.3.2) on gap-free, full-length CDS alignments.

## Results and Discussion

### OA receptors are found in single copy in most of the *Drosophila* genomes we analyzed

To examine the evolutionary history of OA receptors, we first identified the orthologs of OA receptors across 27 *Drosophila* species with available transcriptome data, which, combined with the NCBI Gnomon gene predictions, provides comprehensive and well-supported receptor annotations. Searches against transcriptomes, rather than genomes, are more sensitive to detect paralogs since expressed coding sequences do not have introns that complicate genome-based searches and the database that is being searched against is much smaller. By examining syntenic regions, we found that most OA receptor family members are retained as single-copy genes across species (Fig 1-2, Fig S1-3, File S1), reflecting evolutionary conservation and functional constraint. This is likely important for maintaining a tightly regulated neuromodulatory system, where altered gene dosage or expression levels could lead to signaling imbalances and physiological dysfunction.

**Figure 1.**
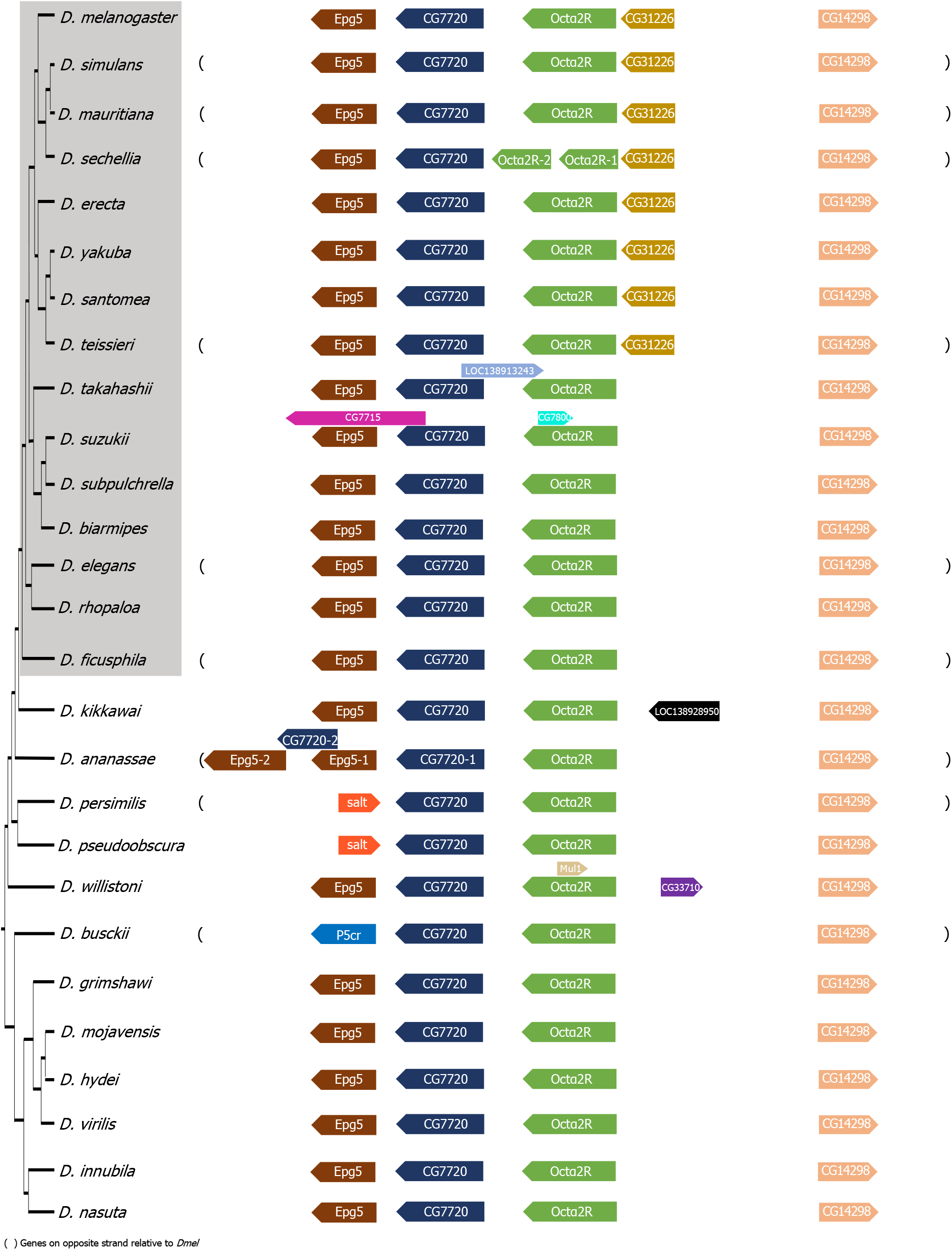
Syntenic region of *Octα2R* across *Drosophila* species. The phylogeny is based on Suvorov et al. (2022) and Hopkins et al. (2024). Surrounding gene names correspond to orthologs in *D. melanogaster*. The species within the gray box represent the *melanogaster* group as defined in the paper.

Interestingly and in contrast, the *Drosophila sechellia* (*Dsec*) genome possesses two copies of *Octα2R* (Fig 1) and *Drosophila busckii* (*Dbus*) genome has two copies of *Octβ3R* (Fig 2). Although the evolutionary forces underlying this variation in OA receptor copy number are unknown, it is intriguing that both *Dsec* and *Dbus* have adapted to thrive on toxic hosts: *Dsec* feeds on *Morinda* fruit (Legal et al., 1992; Louis, 1986), while *Dbus* consumes rotting vegetables such as potatoes (Atkinson, 1977; Buda et al., 2009). This raises the possibility that OA signaling may play a role in adaptation to environmental conditions through lineage-specific diversification or amplification. Since dopamine and octopamine signaling interact functionally in flies (Burke et al., 2012; Ma et al., 2016; Sabandal et al., 2020; Schwaerzel et al., 2003), and the dopaminergic system is crucial for *Dsec*’s reproductive success and specialization on its toxic host (Lavista-Llanos et al., 2014), it is plausible that octopaminergic signaling may also have contributed to such species-specific adaptations. Gene copy number changes have been linked to ecological adaptation in other species. For example, human populations with high-starch diets exhibit increased amylase gene copy number relative to those with lower-starch diets (Perry et al., 2007).Together, these findings raise the possibility that OA receptor copy number variation may contribute to ecological specialization and environmental adaptation. Alternatively, this variation may reflect reduced selective restraint on these genes in these species, leading to toleration of duplication.

**Figure 2.**
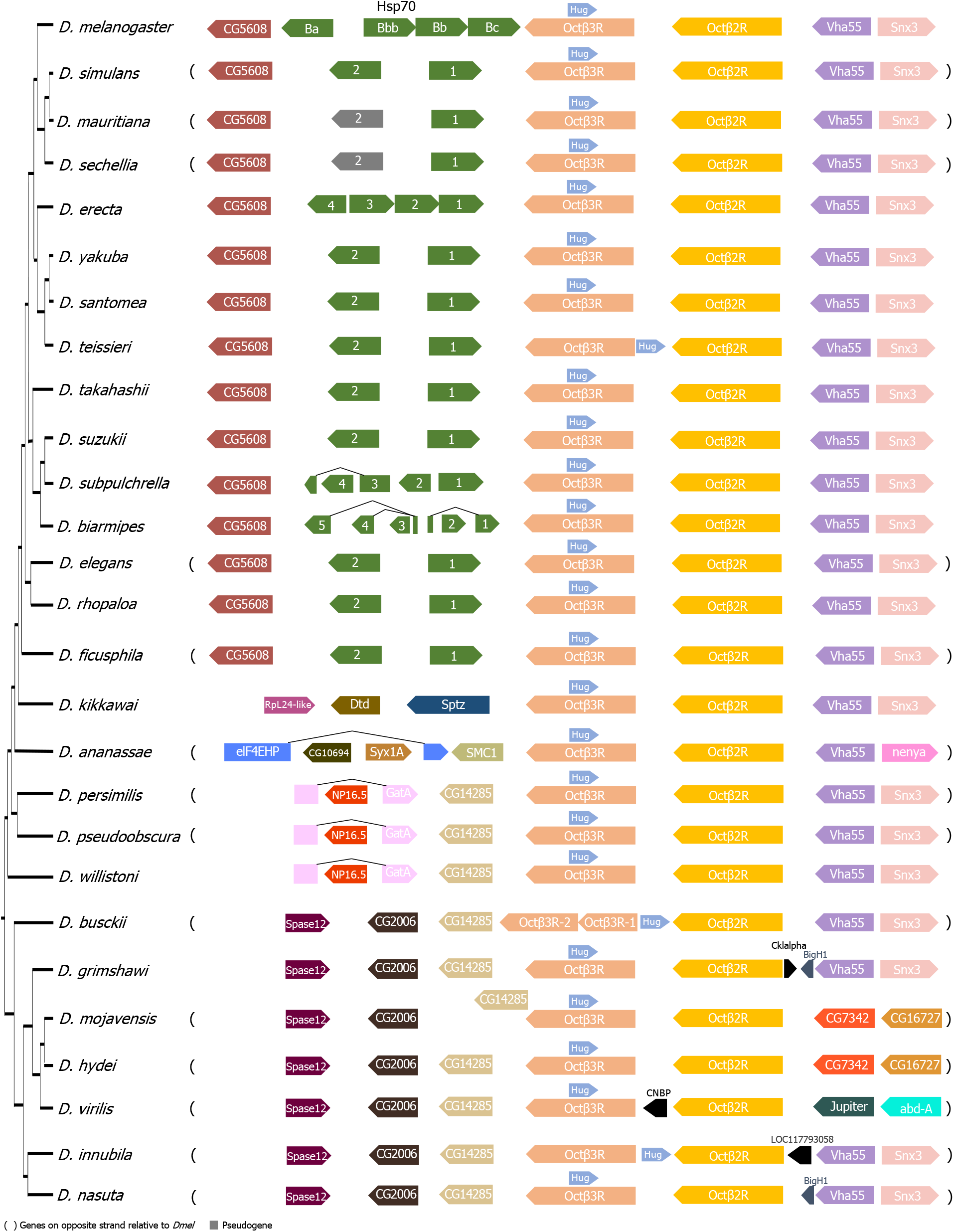
Syntenic region of *Octβ2R* and *Octβ3R* across *Drosophila* species. The phylogeny is based on Suvorov et al. (2022) and Hopkins et al. (2024). Surrounding gene names correspond to orthologs in *D. melanogaster*.

### Octβ2R and Octβ3R have undergone positive selection in the *Drosophila melanogaster* group

We then used PAML to investigate whether any of the OA receptors contain positively selected sites (Yang, 2007), focusing our analysis on species within the *melanogaster* group (Fig 1, S4), which corresponds to Clade 4 in the phylogeny of Suvorov et al. (2022). Because the functions of these genes have been characterized in *D. melanogaster*, limiting our selection analyses to this clade allows for evolutionary insights that are more directly relevant to the experimentally studied functions. Model M0 was used to estimate the overall *d*_N_*/d*_S_ (ω) ratio across the entire protein-coding sequence of each gene; and models M7 vs M8 and M8a vs M8 comparisons (where M7 and M8a are neutral models and M8 is a model that allows for positively selected sites) were used to identify genes evolving under positive selection and identify specific positively selected sites (Swanson et al., 2003; Yang et al., 2000). The M8a null model explicitly includes neutrally evolving sites, and the M8a/M8 comparison is therefore a more sensitive alternative to M7/M8 and less likely to yield false positives. We consider a gene to be under positive selection only if both comparisons are significant. For genes under positive selection, we use the posterior probabilities from the Bayes Empirical Bayes (BEB) test included in the output for codeml’s M8 for identifying sites under positive selection with a posterior probability threshold of 0.9.

Overall, OA receptors exhibit relatively low M0 ω estimates (*d*_N_*/d*_S_ across the entire gene), which may reflect their conserved roles in neuromodulatory and physiological processes. Octβ2R and Octβ3R have the two highest M0 ω estimates among them, and interestingly, we found significant evidence of positive selection acting on several amino acid sites within each of these two receptors (Fig 3A). These genes have multiple isoforms identified and some of these isoforms vary in coding DNA sequence (CDS) which could lead to isoforms having differing results from site selection tests. We performed our analysis on all isoforms for our genes of interest that are sufficiently annotated across species. The results for different isoforms were largely consistent with one another, however, there was a difference in results for Octβ3R isoforms.

**Figure 3.**
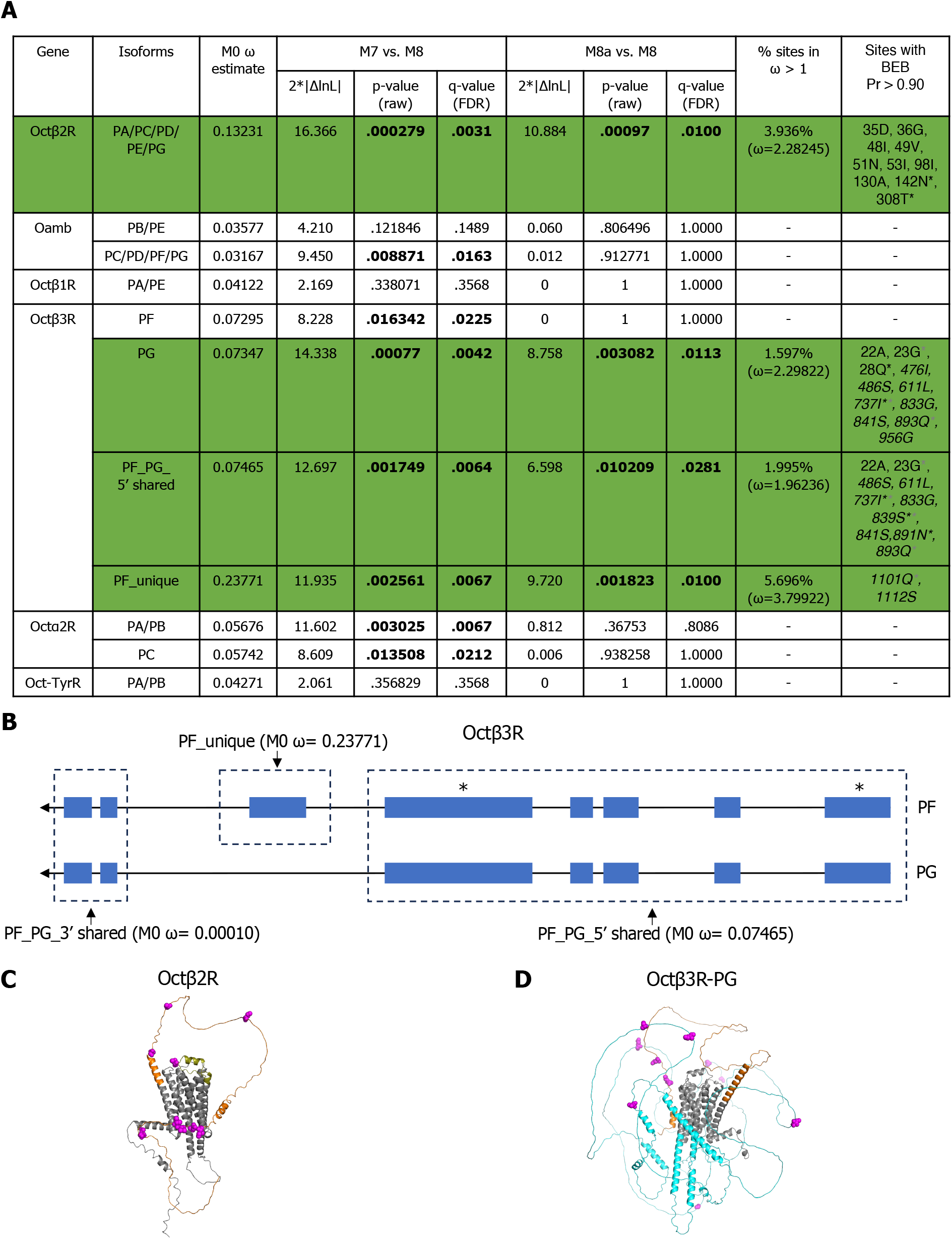
Octβ2R and Octβ3R are under recurrent positive selection in the *melanogaster* group.PAML test results for all six OA receptors across the *melanogaster* group. Significant p-values (raw p-values and Benjamini-Hochberg FDR adjusted p-values (q-values, rounded to four decimal places)) are shown in bold, and coding sequences with significant evidence of positive selection are highlighted in green. The italicized sites in the last column (mapped to *D. melanogaster* amino acid sequence) are located in the intracellular domain, while the others are extracellular. Asterisks in the last column indicate sites also detected as positively selected by MEME (For Octβ3R, gray asterisks indicate sites also detected in the PF isoform MEME analysis, black asterisks indicate sites detected in the PG isoform analysis, and sites marked with both gray and black asterisks were found in both MEME analyses) (File S4, S5). BEB: Bayes empirical Bayes. Schematic of the PF and PG isoforms of Octβ3R. Boxes represent exons, lines indicate introns, and asterisks mark exons containing positively selected sites. The “PF_PG_5’ shared” and “PF_unique” regions tested in panel A, as well as the “PF_PG_3’ shared” exons, are outlined with dotted borders. M0 ω estimations for each region are indicated. C. Positively selected amino acid sites in Octβ2R (BEB posterior probability > 0.90) are shown in magenta and mapped onto the *D. melanogaster* protein sequence and AlphaFold-predicted structure. Nine sites are in the N terminus (orange) and one site is in the extracellular loop 2 (deep olive); both regions are extracellular. D. Positively selected amino acid sites in the PG isoform of Octβ3R (BEB posterior probability > 0.90) are shown in magenta and mapped onto the *D. melanogaster* protein sequence and AlphaFold-predicted structure. Three sites are in the N terminus (orange; extracellular) and eight sites are in the intracellular loop 3 (cyan; intracellular).

Under our criteria for the detection of positive selection, we identified sites under positive selection in the Octβ3R-PG isoform but not the PF isoform (Fig 3A). The only difference in CDS between these isoforms is the presence of an extra exon in the PF isoform (Fig 3B). Notably, the positively selected sites (PSSs) identified in the PG isoform using the BEB posterior probabilities from codeml’s M8 output are all located in regions shared by both isoforms (Fig 3A, 3B). To confirm the reliability of PSS detection, we performed an additional analysis focusing solely on this shared region (PF_PG_5’ shared), which yielded results consistent with those of the PG isoform, showing significant differences in both the M7/M8 and M8a/M8 comparisons, with most PSSs overlapping with those of PG (Fig 3A). Analysis of the PF-unique exon also detected positive selection under both M7/M8 and M8a/M8 comparisons and exhibited the highest M0 ω estimate relative to the shared regions (Fig 3A, 3B). Interestingly, when this exon is included in the full PF isoform, the signal of positive selection is lost. This apparent loss of significance may result from the unstructured nature of intracellular loop 3 encoded by the PF-unique exon, which is likely evolving under reduced constraint. Such variation could alter the underlying ω distribution and reduce the sensitivity of PSS detection in the full isoform. Moreover, the p-values for Octβ2R and Octβ3R-PG remain significant after Benjamini-Hochberg false discovery rate (FDR) correction (q-value; Fig 3A), supporting the robustness of our results.

The PSSs in Octβ2R and Octβ3R are predominantly clustered within unstructured loops (Fig 3C, 3D). In Octβ2R, the PSSs are primarily located in the N terminus, with one additional site in extracellular loop 2 (ECL2) (Fig 3C). Notably, in the human β2-adrenergic receptor, a predicted ortholog of Octβ2R, the N terminus and ECL2 have been reported to be important for ligand binding and accessibility (Isin et al., 2012; Shahane et al., 2014). This suggests that Octβ2R may have evolved to optimize interactions with its ligand. In some *melanogaster* group species, Octβ2R may have evolved to bind OA with altered affinity. Alternatively, since OA is identical across species, this raises the intriguing possibility that Octβ2R could have adapted to bind an additional ligand. In Octβ3R, the PSSs are located in the N terminus and predominantly in intracellular loop 3 (ICL3) (Fig 3D). In class A GPCRs, ICL3 plays a crucial role in signal transduction and receptor activation, with its conformation and length influencing G protein accessibility and selectivity (Sadler et al., 2023). The adaptive evolution of ICL3 in Octβ3R may partially explain why Octβ3R is unable to fully compensate for Octβ2R’s function in ovulation, unlike Octβ1R (Lim et al., 2014). Additionally, Octβ3R may have evolved to interact with different G proteins and/or acquired novel or specialized signaling functions in certain species.

To validate the PAML-based results, we performed complementary analyses, including assessment of mutational bias and additional model-based tests (MEME). To exclude mutational bias as a driver of elevated ω values, we examined codon usage bias, measured by the effective number of codons (ENC), and GC content, including GC3 (GC content in the third codon position), in the *melanogaster* group. Both genes show moderate codon usage bias (ENC: Octβ2R ∼42-50; Octβ3R-PG ∼41-49) and relatively stable GC content across species (Octβ2R: GC ∼0.55-0.58, GC3 ∼0.72-0.81; Octβ3R-PG: GC ∼0.58-0.61, GC3 ∼0.72-0.79) (Files S2), indicating that mutational bias is unlikely to explain the observed patterns. Finally, we used MEME analysis (Murrell et al., 2012) to investigate whether an alternate approach also detected PSSs in Octβ2R and Octβ3R. Unlike PAML, which tests for pervasive selection, MEME detects episodic selection using a different statistical framework. MEME identified more PSSs than PAML, and although only a few sites overlapped between the two analyses (Fig 3A), the PSSs detected by either method were localized in similar regions of the proteins. Detailed discussion of these analyses and supporting data can be found in the supplement (File S3-S5).

In addition, Octα2R and certain isoforms of Oamb also showed sites under positive selection in the M7/M8 comparison but lost significance in the M8a/M8 test (Fig 3A). Since the M8a null model is an adaptation of M7 that includes an additional category of sites where ω = 1 (neutrally evolving), this result suggests that these sites are more likely evolving under neutral rather than adaptive selection. The possibility of redundancy with some of the other OA receptors could also contribute to relaxed selective constraint on Octβ2R and Octβ3R, allowing them to evolve more rapidly.

From a broader perspective, the observation that only some OA receptors have sites that have evolved under positive selection in distinct functional domains highlights the heterogeneous selective pressures shaping the evolution of this receptor family.

### Octβ2R does not contain positively selected sites in the *Drosophila virilis*-*repleta* radiation nor in mosquitoes

We were curious whether positive selection of Octβ2R was restricted to the *melanogaster* clade, or part of a broader pattern of its evolution. To explore this, we examined whether Octβ2R is also under positive selection in other clades. We first analyzed species from the *virilis*-*repleta* radiation,which last shared a common ancestor with the *melanogaster* group approximately 40-60 million years ago within the *Drosophila* genus (Russo et al., 1995; Tamura et al., 2004). Using PAML, we found that both the M7/M8 and more sensitive M8a/M8 comparisons showed no significant difference, suggesting that Octβ2R is under purifying selection in this clade (Fig 4A). To further broaden our analysis, we extended our tests to mosquitoes, a more distantly related group of Dipterans. To minimize the risk of spurious signals of positive selection driven by synonymous site saturation due to excessive evolutionary distance, we adopted a stepwise approach to the PAML analyses. We began with the closely related *Cellia* species, where the M7/M8 and M8a/M8 comparisons revealed no evidence of positive selection (Fig 4A, 4B). We then expanded our analysis to *Anophelinae* and subsequently to the broader *Culicidae* species (Fig 4B), despite the divergence between *Culicinae* and *Anophelinae* exceeding 100 million years (Krzywinski et al., 2006). Across all comparisons, we consistently found no evidence of positive selection (Fig 4A).

**Figure 4.**
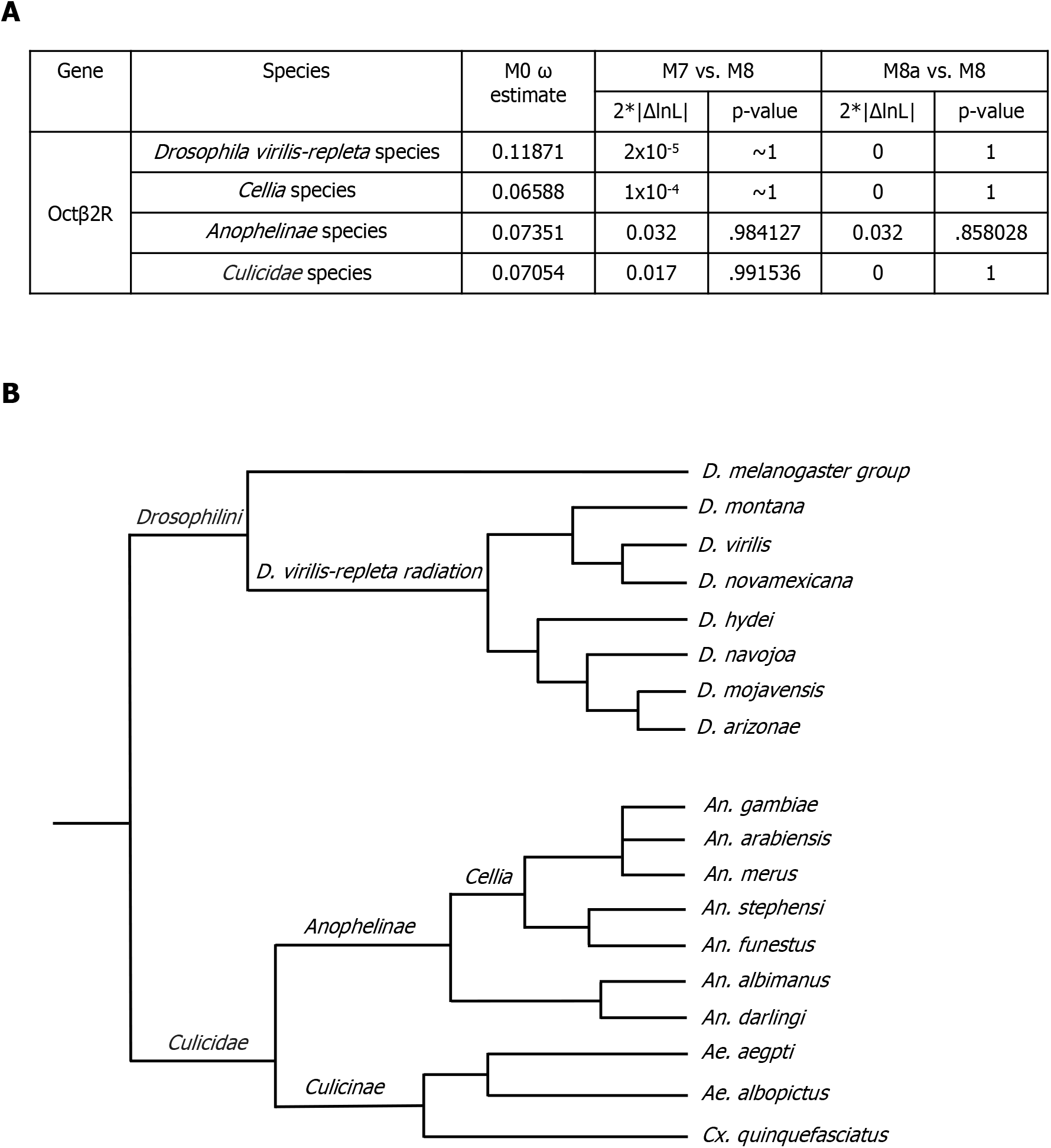
Octβ2R is highly conserved in the *Drosophila virilis*-*repleta* radiation and in mosquitoes. A.PAML test results for Octβ2R across the *virilis*-*repleta* radiation and mosquito (*Cellia, Anophelinae*, and *Culicidae*) species. No significant differences were detected in any M7 vs. M8 or M8a vs. M8 comparisons, indicating that Octβ2R is under purifying selection in these groups. B.Phylogeny of the *virilis*-*repleta* radiation and mosquito species used in the PAML analyses, based on the species tree reported by Suvorov et al. (2022) and Neafsey et al. (2015). The *melanogaster* group is also included to illustrate relationships among the three clades in which Octβ2R was analyzed.

Positively selected sites were detected in Octβ2R within the *Drosophila melanogaster* group, but not in the *Drosophila virilis*-*repleta* radiation or mosquito species, suggesting a lineage-specific change in evolutionary selective pressure. One possibility could be that there is lineage-specific functionality in a process such as reproduction. Previous studies have shown that Octβ2R is essential for ovulation (Li et al., 2015; Lim et al., 2014) and is expressed in both the oviduct epithelium and OA neurons that project into the reproductive tract (Deshpande et al., 2022; Koon et al., 2011). *D. virilis* and *D. melanogaster* vary greatly in the coterie of seminal fluid proteins identified in their genomes and ejaculate (Garlovsky & Ahmed-Braimah, 2023), several of these proteins have been observed to undergo rapid sequence divergence, sometimes driven by positive selection (Haerty et al., 2007; Patlar et al., 2021; Swanson, Clark, et al., 2001; Wong & Wolfner, 2012). Among other hypotheses, it is intriguing to wonder whether rapid evolution of the female reproductive tract-expressed Octβ2R in the *melanogaster* clade might be driven by interaction with a rapidly-evolving ejaculate protein in this clade, such as ovulin, which stimulates ovulation through modulating OA signaling (Aguade, 1998; Aguade et al., 1992; Heifetz et al., 2000; Rubinstein & Wolfner, 2013; Tsaur et al., 1998; Tsaur & Wu, 1997; Wong et al., 2006) and is absent from species like *D. virilis* and *D. mojavensis*. However, experimental validation would be needed to confirm any potential interaction between *D. melanogaster* Octβ2R and ovulin, or any other protein.

## Conclusion

OA receptors are key modulators of physiology and behavior in insects, yet their evolutionary features remain largely unexplored. While their essential roles imply strong functional constraints, their involvement in reproduction-related processes may also subject them to shifting evolutionary selective pressures. In this study, we examined the molecular evolution of OA receptors across *Drosophila* species and uncovered signatures of both conservation and diversification. Most OA receptors were found as single-copy genes, consistent with stringent functional constraints acting on them. However, we identified copy number changes in two species, *Drosophila sechellia* (Octα2R) and *Drosophila busckii* (Octβ3R), which may reflect lineage-specific adaptations. Among the six OA receptors, Octβ2R and Octβ3R within the *Drosophila melanogaster* clade evolve under positive selection and contain positively selected sites in functionally distinct regions of the GPCR domain. This heterogeneity suggests that different OA receptors are subject to distinct selective pressures across the genus *Drosophila* and between paralogs, likely reflecting divergent roles, molecular partners, or regulatory mechanisms. Notably, the detection of positive selection in the ligand-binding involved regions of Octβ2R within the *melanogaster* group, but not in other lineages, raises the possibility of lineage-specific selective pressure on ligand interaction. Together, our findings illustrate the diverse evolutionary trajectories of a closely related receptor family and motivate future functional studies aimed at understanding the molecular and ecological roles of OA receptors in *Drosophila*.

## Data Availability

PAML analysis data, MEME output files and jsons, and PSE files (PyMOL session files) showing the structures of Octβ2R and Octβ3R with annotated PSSs and functional domains (extracellular, transmembrane, and intracellular regions) are provided in the supplementary files.

## Acknowledgments

We thank Drs. Willie Swanson, Geoffrey Findlay, Yasir Ahmed-Braimah, and anonymous reviewers for advice on the selection analyses and/or comments on the manuscript, and members of the Wolfner lab for helpful discussions and suggestions.

