## Supplemental Figure S1 for "Molecular evolution of octopamine receptors in *Drosophila*"

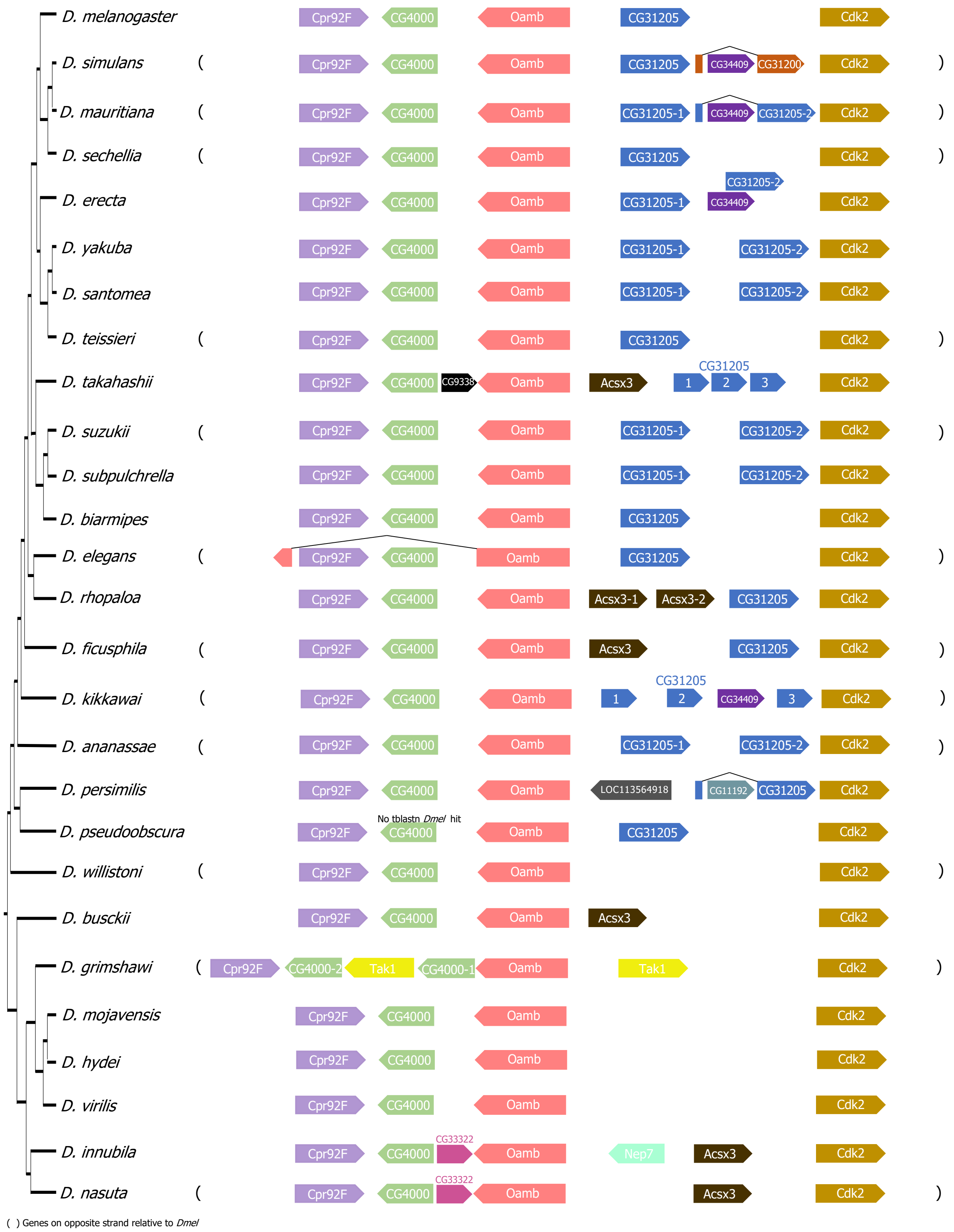


**Figure S1.** Syntenic region of *Oamb* across *Drosophila* species. The phylogeny is based on Suvorov et al. (2022) and Hopkins et al. (2024). Surrounding gene names correspond to orthologs in *D. melanogaster*.
