## Supplemental Figure S4 for "Molecular evolution of octopamine receptors in *Drosophila*"

+: detected -: not detected

| | Octβ2R | Oamb | | Octβ1R | | | Octβ3R | | | | Octα2R | | Oct-TyrR |
|---|---|---|---|---|---|---|---|---|---|---|---|---|---|
| | PA/PC/PD/PE/PG | PB/PE | PC/PD/PF/PG | PA/PE | PB | PC | PF | PG | PJ | PK | PA/PB | PC | PA/PB |
| *D. ficusphila* | + | + | + | + | - | - | + | + | - | - | + | - | + |
| *D. elegans* | + | + | + | + | - | - | + | + | - | - | + | + | + |
| *D. rhopaloa* | + | - | + | + | - | - | + | + | - | - | + | + | + |
| *D. takahashii* | + | + | + | + | + | - | + | + | - | - | + | + | + |
| *D. biarmipes* | + | + | + | + | - | - | + | + | - | - | + | + | + |
| *D. suzukii* | + | + | + | + | - | - | + | + | - | - | + | + | + |
| *D. subpulchrella* | + | + | + | + | + | + | + | + | - | - | + | + | + |
| *D. melanogaster* | + | + | + | + | + | + | + | + | + | + | + | + | + |
| *D. simulans* | + | + | + | + | + | - | + | + | - | - | + | + | + |
| *D. mauritiana* | + | + | + | + | + | - | + | + | - | - | + | + | + |
| *D. sechellia* | + | + | + | + | - | - | + | + | - | - | Duplication | | + |
| *D. santomea* | + | + | + | + | + | - | + | + | - | - | + | + | + |
| *D. yakuba* | + | + | + | + | + | + | + | + | - | - | + | + | + |
| *D. teissieri* | + | + | + | + | - | - | + | + | + | + | + | - | + |
| *D. erecta* | + | + | + | + | - | - | + | + | - | - | + | + | + |

**Figure S4.** Species within the *melanogaster* group used for PAML analyses of OA receptors are shown in gray (+: Orthologs detected by Gnomon prediction. -: Orthologs not detected by Gnomon prediction.). For receptors with multiple different isoforms, only those detected in more than 10 species were included in the analysis. *D. sechellia* was excluded from Octα2R analysis due to the tandem duplication event in this species.
